# Phylogenetic Analysis of Crimean-Congo Hemorrhagic Fever Virus in Inner Mongolia, China

**DOI:** 10.1101/2021.06.06.447231

**Authors:** Yunyi Kong, Chao Yan, Xiaodong Liu, Lingling Jiang, Gang Zhang, Biao He, Yong Li

**Affiliations:** Key Laboratory of Ministry of Education for Protection and Utilization of Special Biological Resources in the Western China, Ningxia University, Yinchuan, 750021, China; School of life science, Ningxia University, Yinchuan, 750021, China; Key Laboratory of Jilin Province for Zoonosis Prevention and Control, Institute of Military Veterinary Medicine, Academy of Military Medical Sciences, Academy of Military Sciences, Changchun, 130062, China

**Keywords:** Inner Mongolia, Crimean Congo hemorrhagic fever virus, phylogenetic analysis

## Abstract

Crimean-Congo hemorrhagic fever virus (CCHFV) is a tick-borne arbovirus that can cause bleeding and death in humans. The mortality rate in humans is between 5 to 30%. The pathogen is prevalent in more than 30 countries in the world. In China, reports of strains of CCHFV have been concentrated in Xinjiang province. However, the CCHFV strain has never been reported in Inner Mongolia, China. This study reports new CCHFV strains, HANM-18, from *Hyalomma asiaticum* and *Hyalomma dromedarii* collected in Alxa Left Banner and Alxa Right Banner in Inner Mongolia. Complete sequences of CCHFV were obtained by the nested PCR technique and used for analyzing the identity and evolutionary relationship with other CCHFV strains. Interestingly, our results showed that the S and L fragments of the HANM-18 strain had high degrees of identity with Xingjiang isolate strains, and the M fragment had significant identity with South African isolates. These analyses also indicate that the HANM-18 strain may have been prevalent in northwest Inner Mongolia for many years. This discovery will be helpful in CCHF prevention and control in Inner Mongolia, and it also adds new evidence to the epidemiology of CCHF in China.

## 1. Introduction

Crimean-Congo hemorrhagic fever virus (CCHFV) is a tick-borne single-stranded RNA virus. The virus belongs to the *Nairoviridae* family of *Bunyavirales* (Garrison et al., 2020). The genome of CCHFV is composed of three fragments (Garrison et al., 2019). The S segment encodes nucleocapsid proteins, the M segment encodes two structural glycoproteins (Gn and Gc), and the L segment encodes an RNA-dependent RNA polymerase (Deyde et al., 2006; Garrison et al., 2019). As for other segmented RNA viruses, rearrangement is also significant in the evolution and migration of CCHFV. Compared with S and L fragments, the rearrangement of the M fragment is more frequent (Deyde et al., 2006). According to phylogenetic analysis, CCHFV can be divided into seven genetic categories related to geographic origin: two from Asia, two from Europe, and three from Africa (Umair et al., 2020).

CCHF caused by CCHFV is a zoonotic disease. It was first reported in Crimea in 1944, and then found in Congo in 1956, which gives it its name (Bente et al., 2013; Spengler et al., 2019). Most infected people are bitten by ticks carrying the pathogen, while others are infected by contact with diseased animals or human body fluids (Garrison et al., 2019; Nabeth et al., 2004; Negredo et al., 2019). The clinical symptoms in humans include fever, headache, vomiting, diarrhea, and muscle pain (Ergönül, 2006). In severe cases, there are signs of bleeding and multiple organ dysfunction (Bente et al., 2013; Ergönül, 2006). Ticks (mainly from the genus *Hyalomma*) are the main vectors of CCHFV; they are not only carriers of the virus, but they are also the natural reservoir of the virus (Spengler et al., 2019; Serretiello et al., 2020). There are no obvious symptoms in mammals infected with CCHFV (Negredo et al., 2019; Spengler et al., 2016). Meanwhile, birds are resistant to CCHFV infection (Negredo et al., 2019). Although some infected animals are usually without infection symptoms, they can still act as hosts in pathogen transmission routes (Serretiello et al., 2020; Spengler et al., 2016; Sorvillo et al., 2020).

The geographical distribution of CCHFV covers most countries and regions in west Asia, the Middle East, Europe, and Africa (Bente et al., 2013; Sorvillo et al., 2020). In China, CCHF infection was first reported in Bachu County of southwest Xinjiang in 1965 (Gao et al., 2010; Wu et al., 2013). Subsequently, there were cases and seropositive reports in the Tarim Basin and surrounding areas (Saijo et al., 2005; Sun et al., 2009). These case reports showed that most of the infected people had close contact with livestock (Gao et al., 2010; Zhang et al., 2018). So far, CCHF cases have only been reported in Xinjiang. However, CCHFV antibodies were detected in livestock and human sera in Qinghai, Inner Mongolia, and Yunnan provinces (Gao et al., 2010; Wu et al., 2013; Xia et al., 2011). This indicates that CCHF may exist in other parts of the Chinese mainland. In previous studies, we reported the detection of CCHFV antibodies in camels and sheep in the Alxa Left Banner and Alxa Right Banner of Inner Mongolia (Li et al., 2020). Nevertheless, the prevalence of CCHFV in this area is not clear. At the same time, the epidemic strain of CCHFV has not been reported in Inner Mongolia.

In this study, we report on the Crimean-Congo hemorrhagic fever virus detected in ticks collected in the Alxa Left Banner and Alxa Right Banner of Inner Mongolia from April to May 2015. The full sequence of the virus was obtained by genome-wide amplification. Phylogenetic analysis and investigation of CCHFV in sheep and camel parasitic ticks were carried out to explore the evolutionary relationship between strains, and the status of local ticks carrying CCHFV was investigated.

## 2. Materials and Methods

### 2.1. Sample collection

On the basis of previous studies, a total of 627 ticks were collected from sheep and camels in the Alxa Left Banner and Alxa Right Banner of Inner Mongolia, China, from April to May 2015 (Figure 1) (Li et al., 2020). For each sampling site, three flocks of sheep or camels were selected, and more than 50 animals in each flock were randomly selected for sampling. Ticks found on the skin surface of the animals were removed and collected in a container. The ticks were initially identified to species by morphological characteristics, and then the species were confirmed based on PCR amplification and sequencing of the cytochrome c oxidase subunit 1 gene (Zahler and Gothe, 1997; Apanaskevich and Horak, 2010; Gou et al., 2018). All samples were frozen and stored at −80 °C. The collected ticks were pooled according to the hosts they were collected from and the tick species . Herdsmen were orally informed about the aims and process of this study. All animal owners agreed and allowed us to collect ticks from their camels or sheep.

**Figure 1.**
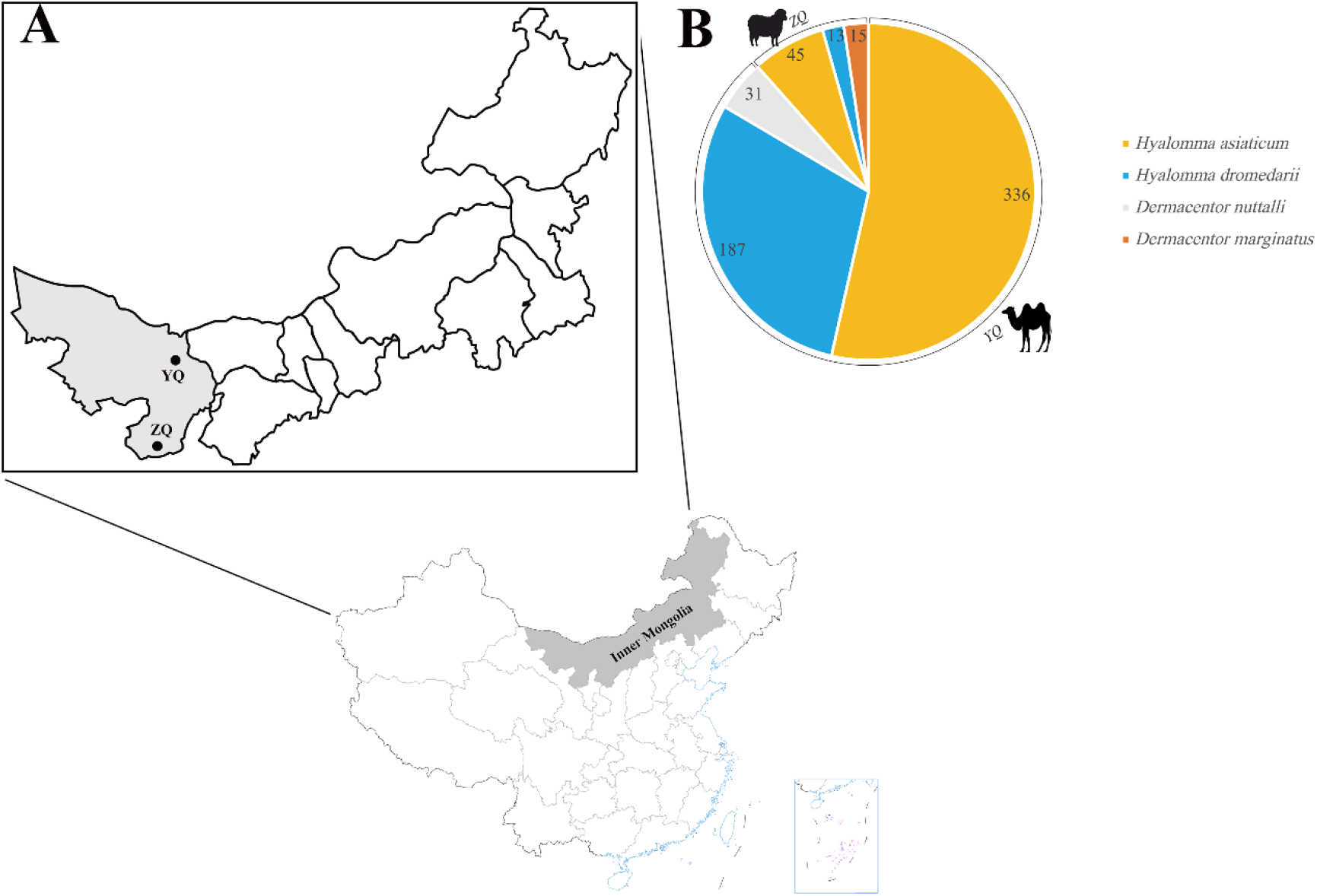
Information for ticks included in the study. (**A**) Sampling locations; (**B**) tick species composition, quantity, and host animals. ZQ: Alxa Left Banner, YQ: Alxa Right Banner.

### 2.2. Sample Processing and RNA Extraction

12-15 ticks were pooled in one sample based on the species and life stage for processing and analysis. To prepare samples, ticks from each pool were put into a 2 mL centrifuge tube containing 1 mL of Dulbecco’s modified Eagle’s medium (DMEM; Gibco, Carlsbad, CA, USA) prior to being homogenized in a frozen grinder (Jingxin, Shanghai, China). The homogenized samples were then centrifuged at 4 °C and 12,000 × g for 10 min. The supernatants were collected and stored at −80°C. Total RNA was extracted from each tick pool with the RNeasy Mini Kit (QIAGEN, Hiden, Germany). According to the manufacturer’s instructions, cDNA was synthesized using M-MLV reverse transcriptase (TaKaRa Biotechnology, Dalian, China). In brief, 0.5 μL dNTP mixture, 1 μL Oligo (dT) 18 primer, 1 μL template RNA (100 ng/μL), and 5 μL RNase-free water were mixed and incubated at 65 °C for 5 min. Next, 2 μL 5x reverse transcriptase M-MLV Buffer, 0.25 μL RNase inhibitor (40 U/μL), and reverse transcriptase M-MLV (200 U/μL) were added to the tube, sequentially incubated at 42 °C for 1 h, and then at 70 °C for 15 min. The cDNA was then stored at −40 °C.

### 2.3. CCHFV detection and whole-genome amplification

To detect whether CCHFV was present in the tick pools, part of the S fragment was amplified by specific nested PCR for detection. The PCR reaction system was a 50 μL mixture, including 25 μL 2x Taq PCR master mix, 2 μL template cDNA, 19 μL ddH_2_O and 20 pmol of each forward and reverse primer. For the first round of PCR, a pair of primers (CCHFV-F 5’-TGGACACYTTCACAAACTC-3’, CCHFV-R 5’-GRYRAAYTCCCTRCACCA-3’) was used to amplify a 536 bp fragment. The PCR amplification was performed in a power cycler (Analytikjena, Jena, Germany) using the following procedure: pre-denaturation at 94 °C for 3 min, then denaturation at 94 °C for 30 s, annealing at 54 °C for 40 s, and extension at 72 °C for 40 s. This cycle was repeated for a total of 35 cycles, followed finally by maintaining at 72 °C for 7 min. For the second round of PCR, another pair of primers (CCHFV-F’ 5’-GAATGTGCATGGGTTAGCTC-3’, CCHFV-R’ 5’-GACATCACAATTTCACCAGG-3’) was used to amplify a 260 bp fragment. The amplification was performed using the following program: pre-denaturation at 94 °C for 3 min, denaturation at 94 °C for 30 s, annealing at 54 °C for 40 s and extension at 72 °C for 30 s. This cycle was repeated for a total of 35 cycles, followed finally by holding at 72 °C for 7 min. The PCR products were analyzed by agarose nucleic acid gel electrophoresis and sent to ComateBio (Changchun, China) for Sanger sequencing.

Genomic amplification was carried out in tick pools where CCHFV was detected. The semi-nested PCR primers (Table S1) were designed to amplify the whole genome sequence of CCHFV. The full sequences were amplified by LA Taq DNA polymerase (TaKaRa Biotechnology, Dalian, China) under the following conditions: pre-denaturation at 94 °C for 1 min, then denaturation at 98 °C for 10 s, annealing at 55 °C for 30 s, and extension at 72 °C for 30-90 s. This cycle was repeated for a total of 35 cycles, followed finally by holding at 72 °C for 10 min.

### 2.4. Phylogenetic analysis

Viral sequences were assembled and aligned using Clustal W and compared with the sequences of other CCHFV strains in GenBank (Thompson et al., 2003). Respectively, the best-fit models (the Kimura 2-parameter model and the JTT matrix-based model) of nucleotide and amino acid sequence evolution were estimated using MEGA 7.0 (Kimura 1980; Jones et al., 1992; Kumar et al., 2016). Phylogenetic trees were constructed using the maximum-likelihood method by MEGA 7.0 software (http://www.megasoftware.net/). The reliability of the branches of the tree was assessed using a bootstrap analysis with 1000 replicates. The phylogenetic trees were edited and visualized with FigTree v.1.4.4 (http://tree.bio.ed.ac.uk/software/figtree). The MegAlign program implemented in the Lasergene software package v5.0 (DNAstar, Madison, WI, USA) was used to calculate the identity between nucleic acid and amino acid sequences.

### 2.5. Data analysis

The positive rate was determined by calculating the minimum infection rate (MIR), which was the ratio of the number of positive tick pools to the total number of analyzed ticks (Walter et al., 1980).

## 3. Results

### 3.1. Detection of CCHFV in Ticks

According to the tick species, sampling site, and host animal species, 73 ticks (divided into five tick pools) were collected in Alxa Left Banner, and 554 ticks (divided into 36 tick pools) were collected in Alxa Right Banner (Table 1). Nested PCR was used to test for presence of CCHFV in these 41 tick pools. By PCR amplification, CCHFV was detected in only one tick pool in Alxa Left Banner, consisting of *Hyalomma asiaticum*. The minimum infection rate (MIR) of CCHFV was 2.2% in Alxa Left Banner. Meanwhile, CCHFV was detected in one-sixth of the tick pools in Alxa Right Banner, including four *H. asiaticum* pools and two *Hyalomma dromedarii* pools. In this area, the MIRs of CCHFV were 1.2% and 1.1%, respectively (Table 1). CCHFV was not detected in *Dermacentor marginatus* from Alxa Left Banner or *Dermacentor nuttalli* from Alxa Right Banner.

**Table 1.** Molecular detection of CCHFV from ticks isolated from different regions in Inner Mongolia, China

| Areas of Sampling | Host species | Tick species | Tick life stage | Number of ticks | Number of pools | Number of positive pools | Positive rate (%) |
| --- | --- | --- | --- | --- | --- | --- | --- |
| Alxa Left Banner | Sheep | *Hyalomma asiaticum* | Adult (Female) | 45 | 3 | 1 | 2.2 |
|  |  | *Hyalomma dromedarii* | Adult (Male) | 13 | 1 | 0 | 0 |
|  |  | *Dermacentor marginatus* | Nymph | 15 | 1 | 0 | 0 |
| Alxa Right Banner | Camel | *Hyalomma asiaticum* | Adult (Female) | 336 | 22 | 4 | 1.2 |
|  |  | *Hyalomma dromedarii* | Adult (Female) | 187 | 12 | 2 | 1.1 |
|  |  | *Dermacentor nuttalli* | Nymph | 31 | 2 | 0 | 0 |

### 3.2. Homology and Phylogeny Analysis

The new strain of CCHFV genome which was a genetic sequence recovered from tick pools, named HANM-18, consisted of three fragments: S (1,672 nt), M (5,368 nt), and L (12,155 nt) (Genbank Accession Numbers: MN832721–MN832723). Phylogenetic analysis of the HANM-18 strain showed that there were three complete sequences.

#### 3.2.1 S fragment phylogenetic analysis

According to their geographical distribution, S fragments were divided into eight genotypes, namely Asia 1 and 2, Africa 1–3, and Europe 1–3 (Guo et al., 2017). The typing of M and L fragments was described previously (Deyde et al., 2006; Morikawa et al., 2007; Carroll et al., 2010). Most of the previously reported Chinese strains were clustered in the Asia 2 group, while WJQ16206 and FK16116 were clustered in Asia 1 (Zhang et al., 2018). In this study, HANM-18 and Chinese isolates were clustered into the Asia 2 group. According to Figure 2A, the evolutionary relationship between HANM-18 and the currently reported Chinese isolates shows that they have the same ancestral strain, while the evolutionary relationship between HANM-18 and the ancestral strains seems to be closer. In the homologous relationship, table S2 shows that the S fragment of the HANM-18 strain in this study had the highest identity with the CCHFV isolate in Xinjiang, China, ranging from 95.5–96.6%. Among these strains, HANM-18 had the highest nucleic acid identity with CCHFV discovered in Xinjiang in the 1960s and 1970s.

**Figure 2.**
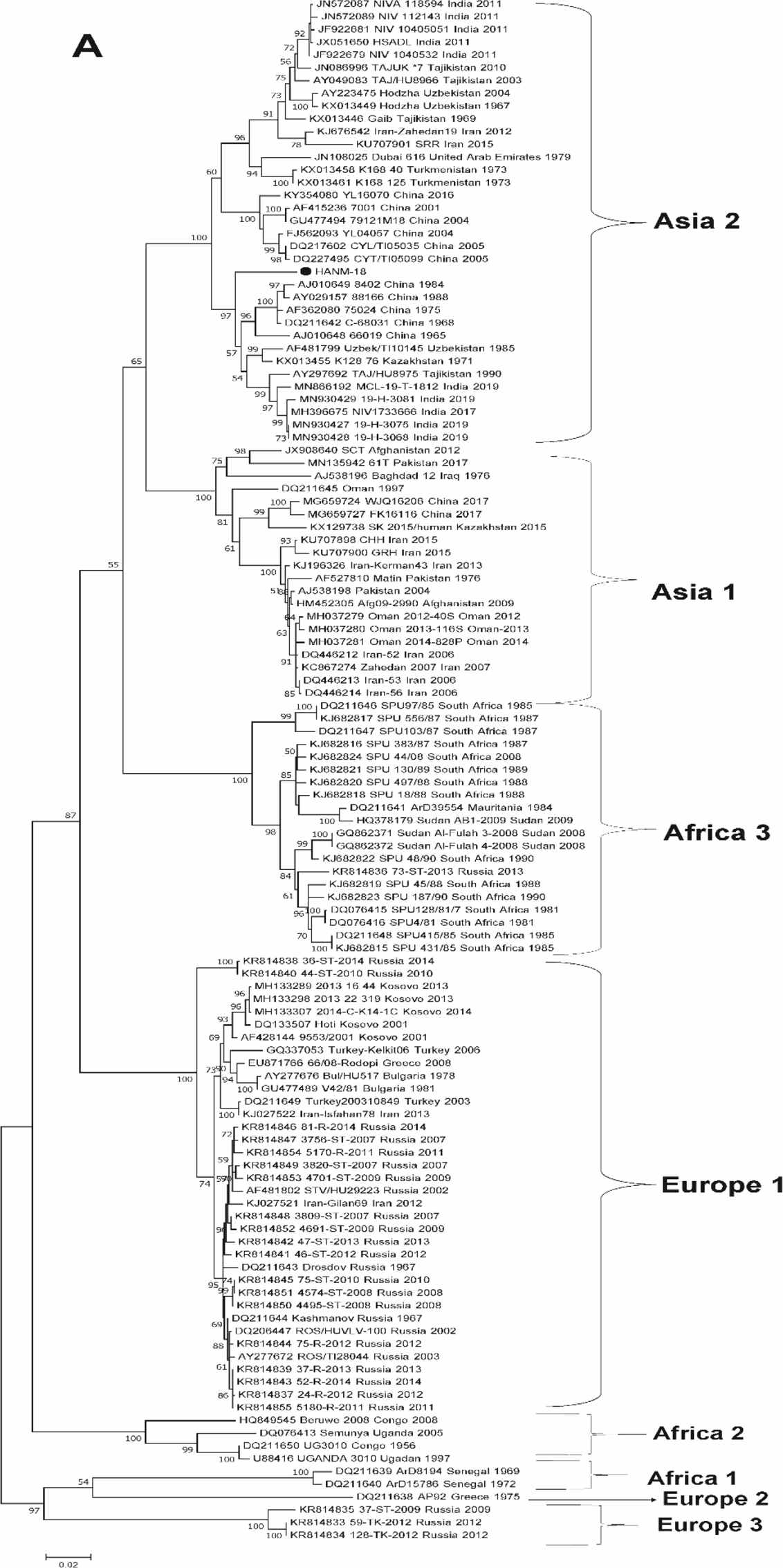
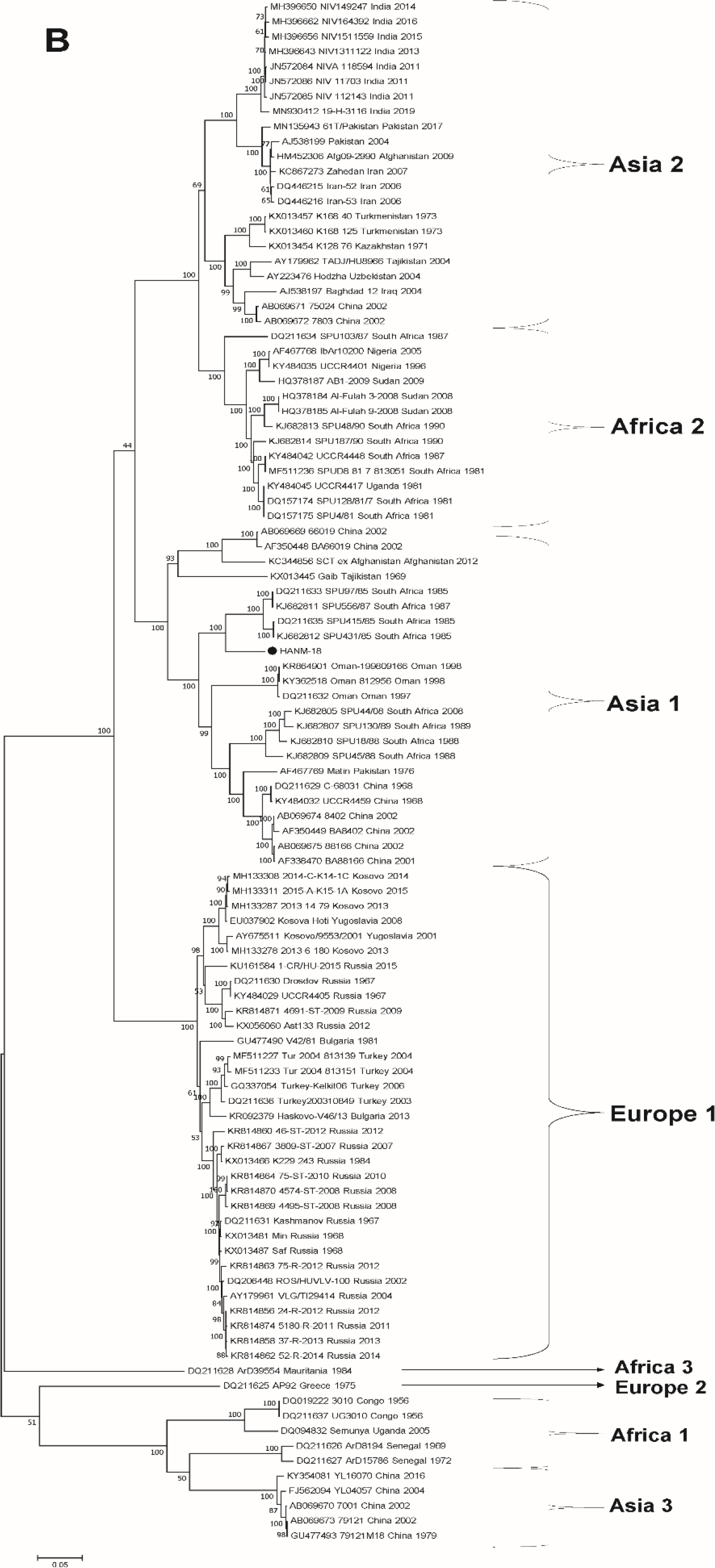
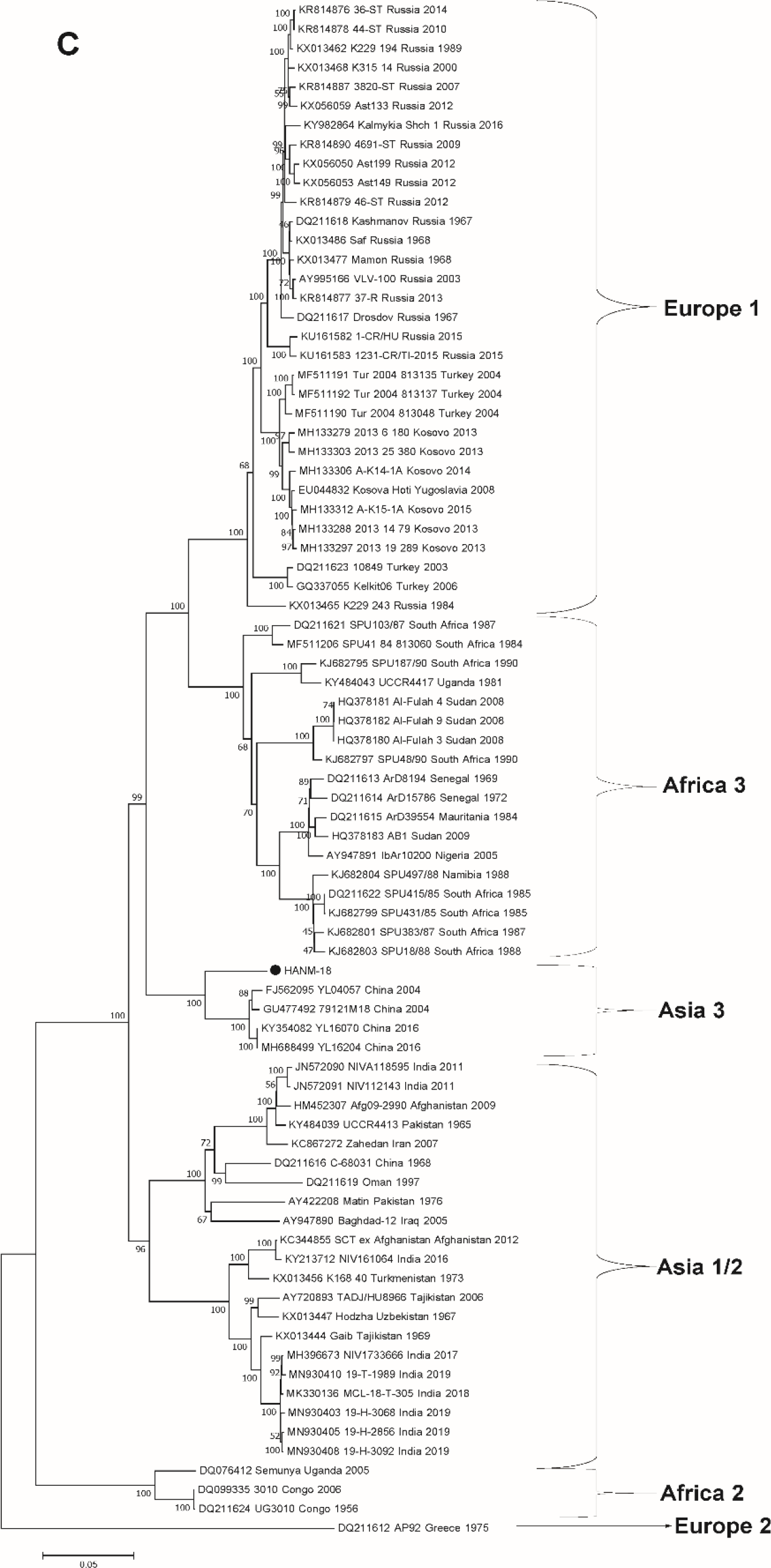
Maximum likelihood trees of Crimean-Congo hemorrhagic fever virus strains based on the complete sequences of the S (**A**), M (**B**), and L (**C**) segments. The solid triangles represent the same type of strains, and the hollow triangles represent the strains of the same country. The size of the triangle has nothing to do with the number of substituting strains. The new strain in this study is highlighted with a black solid circle.

#### 3.2.2 M fragment phylogenetic analysis

As for the M tree, it showed a different evolution from the currently reported Chinese strains. The M fragment of HANM-18 was clustered with an isolate from South Africa (Figure 2B). The Chinese CCHFV strains demonstrated different aggregation phenomena on the phylogenetic tree. The identity between the HANM-18 strain and South African strains (SPU97/85, SPU415/85, and SPU556/87) was 91.1% (Table S3). The homology of HANM-18 with South African strains such as SPU103/87, SPU187/90, SPU48/90, SPU130/89, SPU44/08, and SPU431/85 ranged from 80.8% -90.8%. The identity with some strains (8402, 88166, BA8402, BA88166, and C-68031) from Xinjiang in China was 87%, and the identity with the Oman strain was 86%. Compared with the amino acid sequence identity of the M fragment, it was observed that the amino acid identity of HANM-18 and South Africa was the highest, which was 91.8–91.9% (Table S3). Simultaneously, the identity with some Chinese isolates, Oman strains, and Pakistani strains was over 89%. To further determine the evolutionary relationship of the M fragment in the HANM-18 strain, the sequences with more than 85% amino acid identity were selected to construct the ML phylogenetic tree. In this amino acid tree, the HANM-18 strain was also clustered with South African isolates, and HANM-18 had a distant genetic relationship with Chinese isolates (Figure 3).

**Figure 3.**
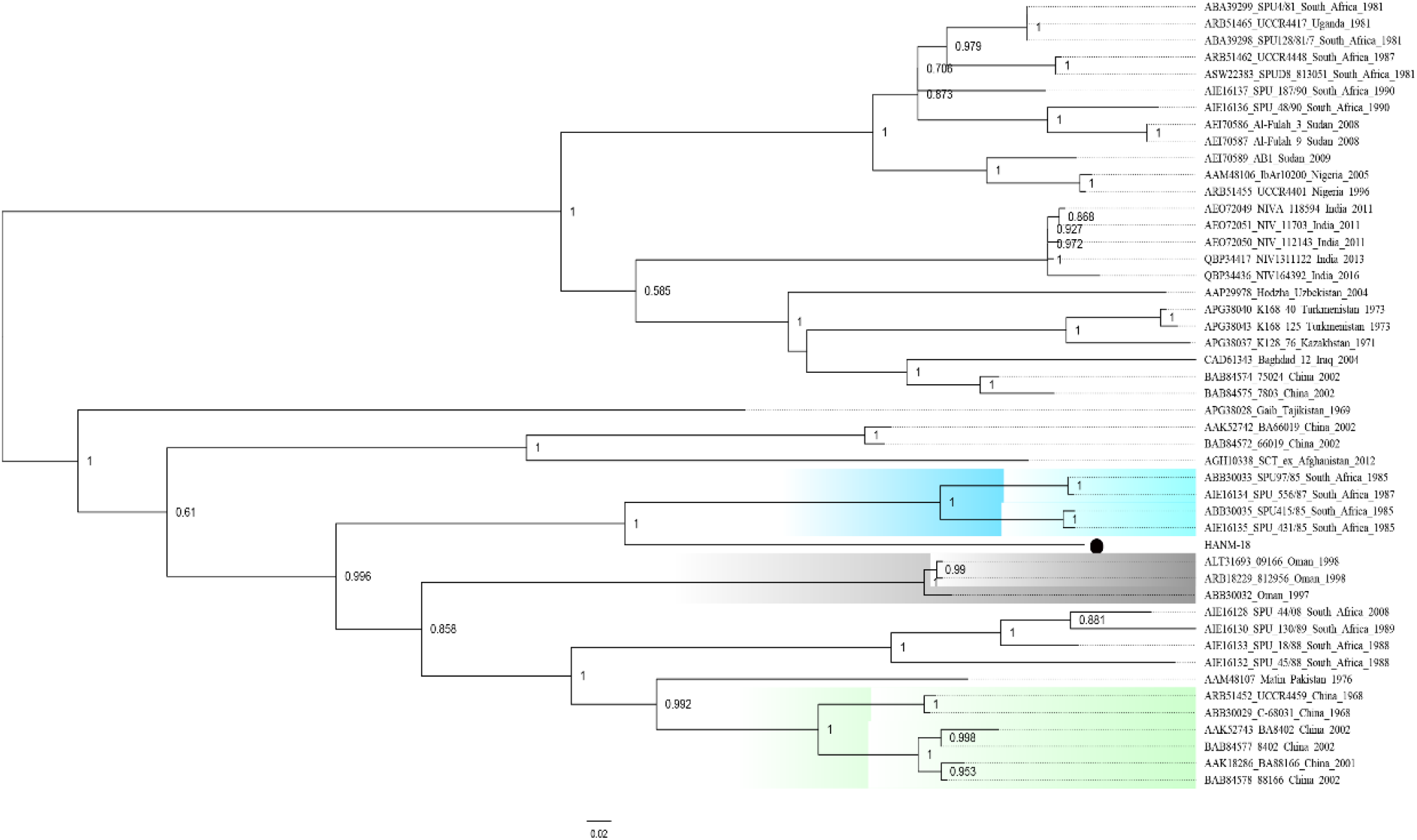
A phylogenetic tree for Crimean-Congo hemorrhagic fever virus based on the glycoprotein precursor was constructed using the maximum likelihood method. The blue, gray, and green areas are used to represent the strains from South Africa, Oman, and China. The new strain in this study is highlighted with a black solid circle.

#### 3.2.3 L fragment phylogenetic analysis

In the L tree, the strain was clustered with YL04057 and 79121M18, discovered in China in 2004, and YL16070 and YL16204 discovered in 2016 (Figure 2C). However, it was far away from the C-68031 strain discovered in 1968. The homology of HANM-18 with YL04057, 79121M18, YL16070, and YL16204 was the highest (93.7%), while that with c-68031 was 87.5% (table S3).

## 4. Discussion

Crimean-Congo hemorrhagic fever virus is a tick-borne virus in many countries in Asia, Africa, and Europe. In China, CCHFV is mainly distributed in Xinjiang province, and no cases have been reported in other provinces. In order to investigate the distribution of CCHFV in northwestern China, ticks were collected and tested in the western region of Inner Mongolia adjacent to Xinjiang.

In the current study, all ticks found to carry CCHFV were of the *Hyalomma* genus. *Hyalomma asiaticum* has been confirmed to be the main vector of CCHFV in Xinjiang, China (Guo et al., 2017; Moming et al., 2018). *Hyalomma dromedarii*, as a tick species mainly parasitic to camels, has also been demonstrated to carry CCHFV and infect the host (Camp et al., 2020; Moshaverinia and Moghaddas 2015). Strangely enough, *H. dromedarii* was found in both Left and Right Banner, while CCHFV was detected only in *H. dromedarii* of females from Alxa Right Banner. In the previous CCHFV survey in Saudi Arabia, CCHFV was also detected in *H. dromedarii* of females (Camp et al., 2020).

The identity and phylogeny analysis of CCHFV demonstrated that CCHFV, as a segmented RNA virus, showed high genetic diversity (Deyde et al., 2006). In the phylogenetic tree of S fragments, the isolates of HANM-18 from Xinjiang were grouped into Asia 2, and China is the origin of Asia 2 (Figure 2A) (Mild et al., 2010). And the homology with the previously reported strains in Xinjiang were the highest (Table S2).For the L fragment, the homology of HANM-18 with some Xinjiang strains (YL04057, 79121M18, YL16070, and YL16204) were the highest. HANM-18 was clustered with the Xinjiang strains in Asia 3 group on the L tree (Figure 2C). It is far away from some strains found in the Middle East. For the M fragment, HANM-18 was clustered with strains from South Africa (SPU97/85, SPU415/85, SPU556/87, and SPU556/87), but is far away from Chinese isolates (Figure 2B,). The identity between HANM-18 and South African strains is also the highest. The amino acid tree of the M fragment and amino acid identity analysis also reached the same conclusion (Figure 3).

For CCHFV, due to its wide geographical distribution, there are significant differences among different epidemic areas, and the variation in different fragments is also dissimilar: from 10–20% for the S segment to 31% of the M segment (Deyde et al., 2006; Bente et al., 2013; Carroll et al., 2010). In general, the S segment is the slowest and most conserved, followed by the L segment, while the M segment shows great differences (Bente et al., 2013; Morikawa et al., 2007). The M fragment of HANM-18 is highly homologous with the South African isolate and is closer to the ancestor of the South African isolate in the genetic relationship (Figure S1B). This suggests that the M fragment of HANM-18 may be from an older strain. Then, its geographical isolation in China was characterized, and adaptive evolution occurred in the process of transmission. There are big differences compared with the strains prevalent in South Africa. At the same time, HANM-18 was far away from the Xinjiang strain in the phylogenetic tree (Figure S1B). This indicates that there are different epidemic types of the CCHFV M fragment in China. The reason for this phenomenon is not only mutation, but also rearrangement between different strains (Varsani et al., 2018). The S, M, and L fragments of CCHFV showed different prevalence and aggregation in China, which indicated that rearrangement was also common and played an important role in the transmission and evolution of CCHFV (Zhou et al., 2013). In previous reports, the South African isolate spu415/85 was closely related to Asian rearrangement (Goedhals et al., 2014). The strains found in South Africa have been confirmed to be prevalent and distributed in China (Zhou et al., 2013; Goedhals et al., 2014). There are many reasons for these rearrangements, either because migratory birds have carried the virus to different places, or because of the trade activities of livestock; ticks parasitized on these livestock have been taken to all parts of the world (Deyde et al., 2006; Serretiello et al., 2020; Spengler et al., 2016; Mild et al., 2010; Zhou et al., 2013; Palomar et al., 2013; Mancuso et al., 2019). In addition, the evolutionary relationship of the M fragment in the CCHFV strain found in Mongolia showed the same phenomenon as that of HANM-18 during the last 13–14 years (Voorhees et al., 2018). Inner Mongolia borders Mongolia, and it is possible that the CCHFV strain will spread between the two regions with the migration of birds or wild animals. However, since the data have not been published, it is impossible to make more comparisons.

In summary, this study indicates the presence of CCHFV strains in Inner Mongolia in China. The genetic relationship is not only related to the strains isolated from Xinjiang, but also shows that there are different types of CCHFV in China. This is the first time that CCHFV strains have been found and reported outside Xinjiang in China, which provides a good reference to characterize the prevalence and migration of CCHFV in China and Asia.

## 4. Conclusions

The present study provides data on the prevalence of important zoonotic agents in ticks from sheep and camels in Inner Mongolia of China. This novel information will be useful not only for medical and veterinary practitioners, but also the public health officials, when assessing the risks associated with tick-borne zoonoses in China. In addition, the authors hope that this report will also contribute, to increasing an overall awareness of tick-borne diseases among the population of China.

## CRediT authorship contribution statement

**Yunyi Kong:** Methodology, Investigation, Data curation, Formal analysis, Writing-original draft. **Chao Yan:** Methodology, Investigation, Data curation. **Dongxiao Liu:** Investigation, Data curation. **Lingling Jiang:** Methodology, Investigation. **Gang Zhang:** Investigation, Data curation. **Biao He:** Formal analysis, Resources. **Yong Li:** Project administration, Conceptualization, Supervision, Writing - review & editing.

## Acknowledgements

This work was supported by Ningxia University (China) and the Academy of Military Medical Sciences (China). We thank Zhizhou Tan for his expert technical assistance.

## Animal and Human Right Statement

The experiments involving sheep and camels were performed according to protocols approved by the Institutional Animal Care and Use Committee of Ningxia University (NXU-2019-007). The need for consent is deemed unnecessary according to national regulations, but an informed verbal consent was obtained from the sheep and camel owners. The ethics committee for the use of animals of Ningxia University approved this study.

## Appendix A. Supplementary data

Table S1-S4

---

**Table S1.**
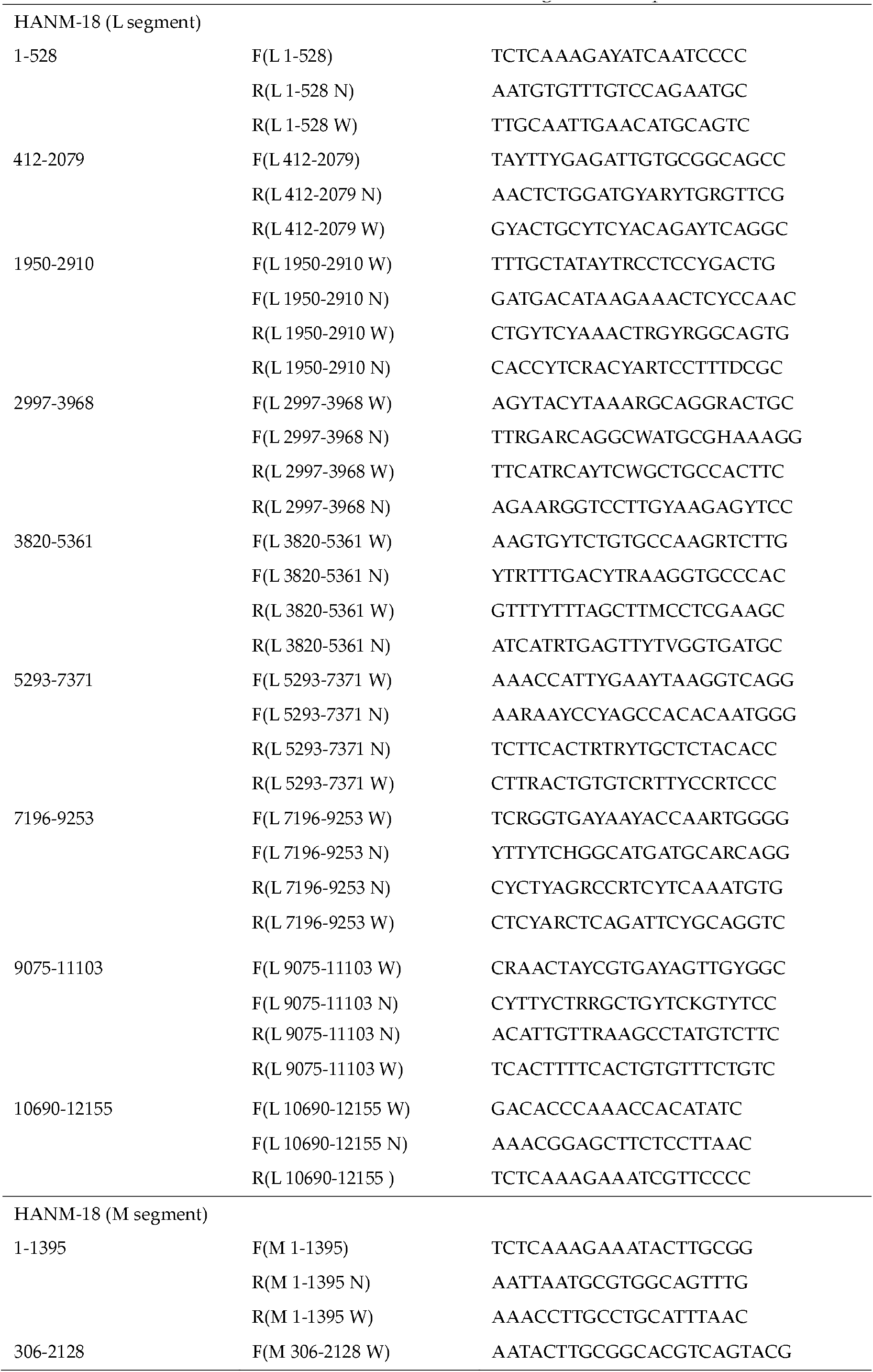
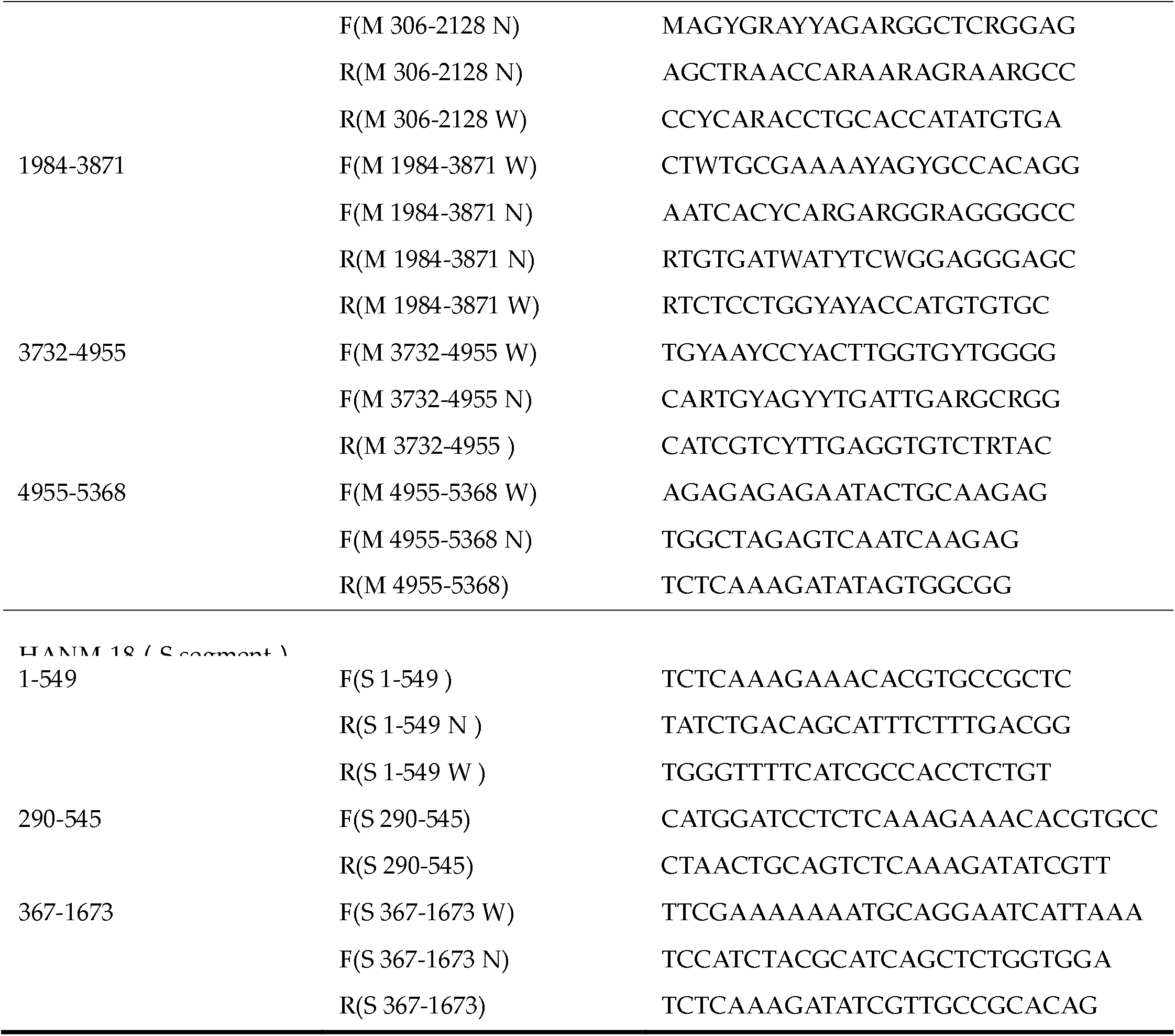
Primer used in the CCHFV whole-genome amplification

**Table S2.**
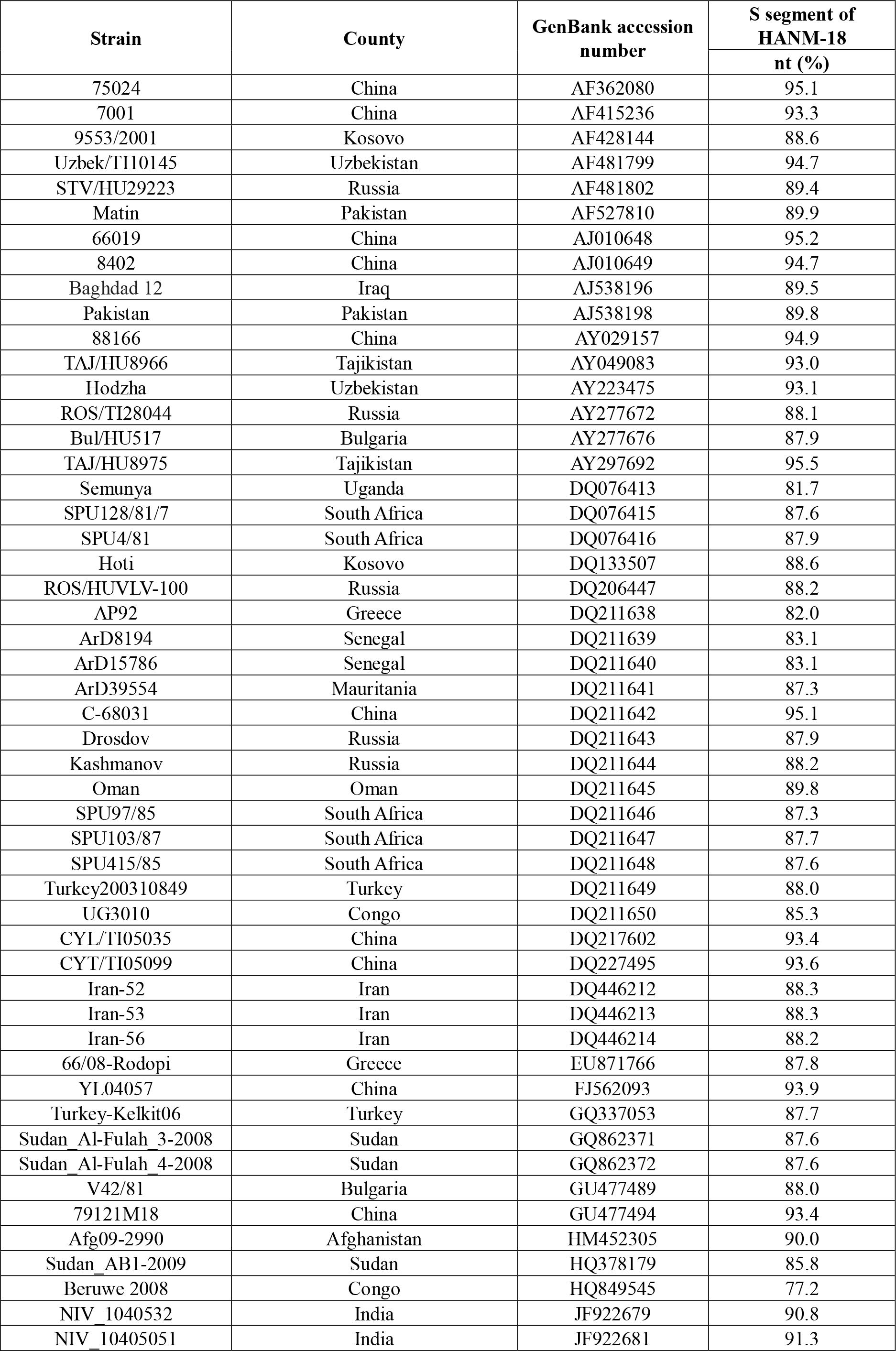
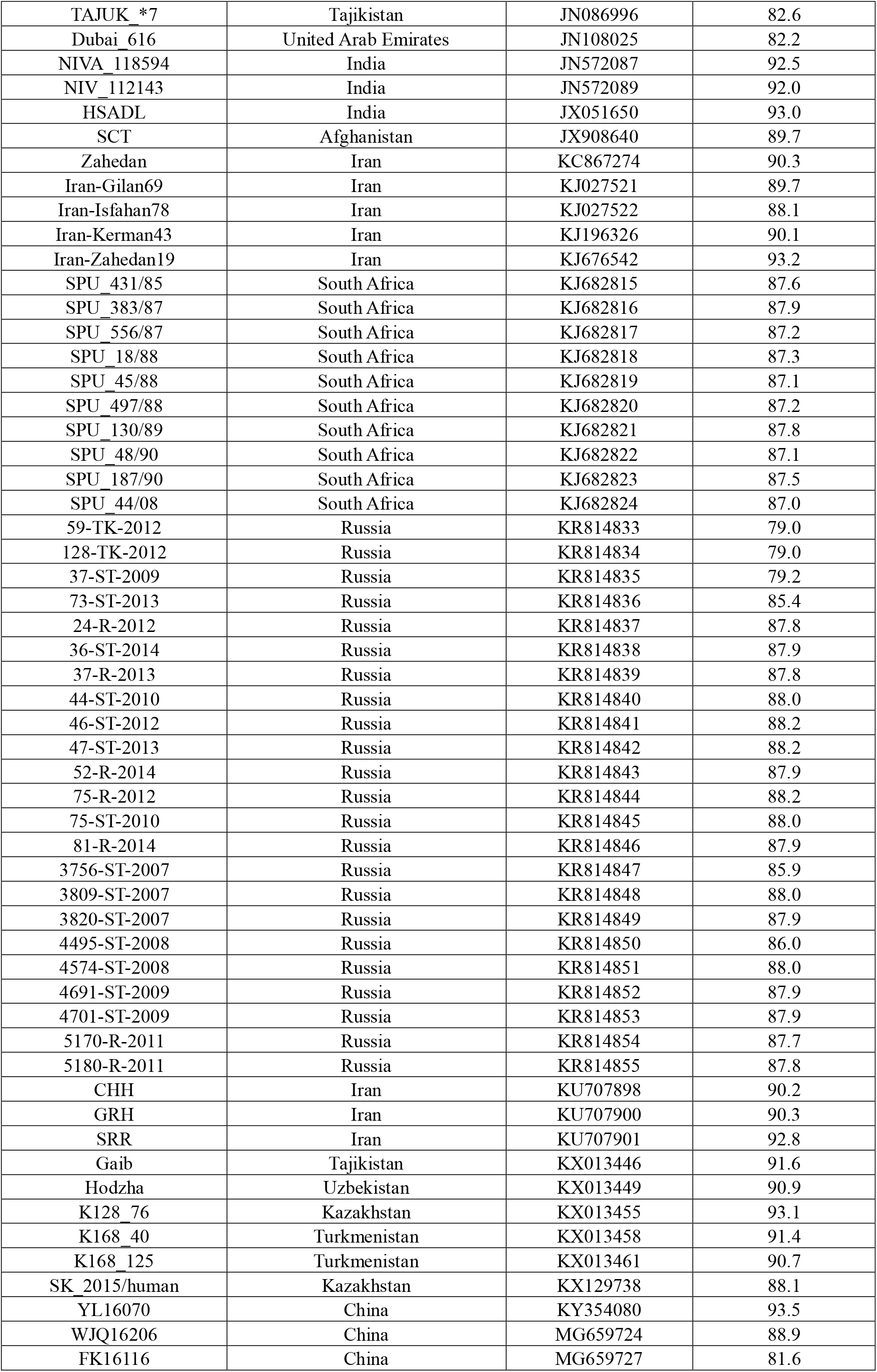
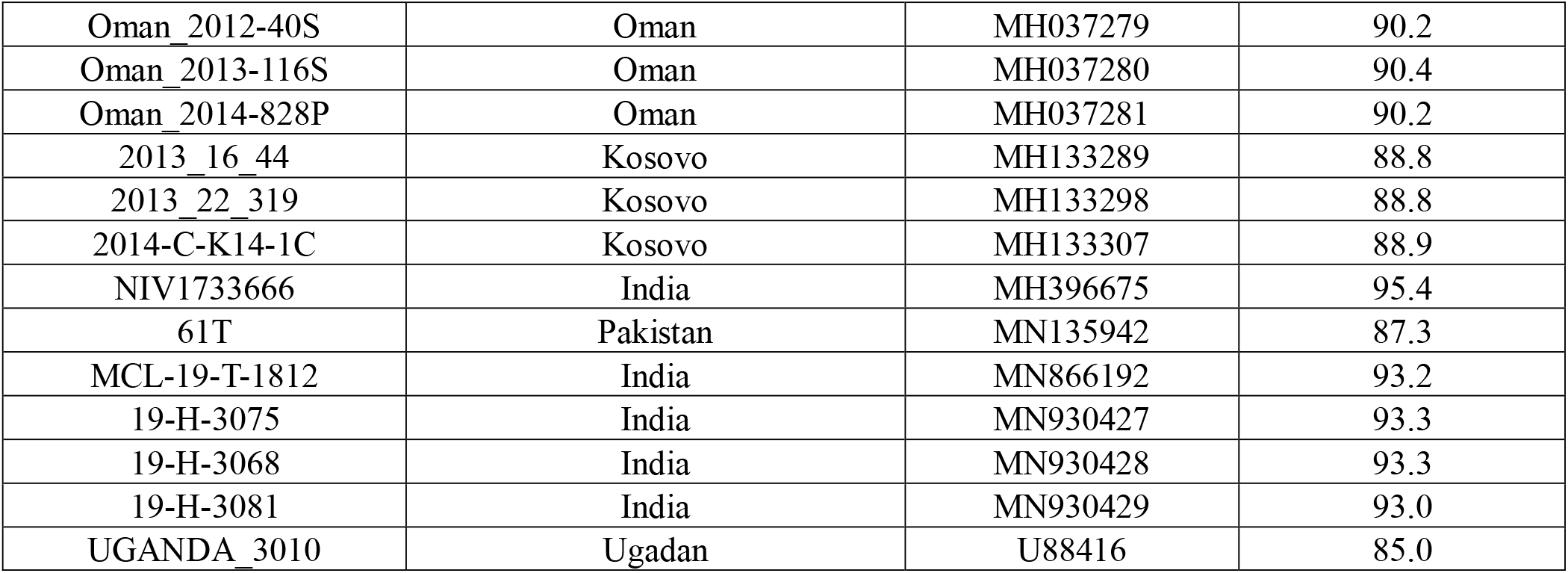
Comparison of nucleotide sequence identity with other CCHFV strains’ S segment

**Table S3.** Comparison of amino acid and nucleotide sequence identity with other CCHFV strains’ M segment

| Strain | Country | GenBank accession number | M segment of HANM-18 |  |
| --- | --- | --- | --- | --- |
|  |  |  | nt (%) | aa (%) |
| 66019 | China | AB069669 | 84.4 | 87.4 |
| 7001 | China | AB069670 | 69.8 | 72.6 |
| 75024 | China | AB069671 | 81.2 | 85.7 |
| 7803 | China | AB069672 | 80.9 | 85.5 |
| 79121 | China | AB069673 | 69.7 | 72.3 |
| 8402 | China | AB069674 | 87.5 | 89.3 |
| 88166 | China | AB069675 | 87.5 | 89.4 |
| BA88166 | China | AF338470 | 87.4 | 89.4 |
| BA66019 | China | AF350448 | 84.3 | 87.4 |
| BA8402 | China | AF350449 | 87.2 | 88.7 |
| IbAr10200 | Nigeria | AF467768 | 81.5 | 85.1 |
| Matin | Pakistan | AF467769 | 87.8 | 90.1 |
| Baghdad 12 | Iraq | AJ538197 | 81.0 | 85.8 |
| Pakistan | Pakistan | AJ538199 | 80.5 | 84.1 |
| VLG/TI29414 | Russia | AY179961 | 80.5 | 84.2 |
| TADJ/HU8966 | Tajikistan | AY179962 | 80.0 | 84.4 |
| Hodzha | Uzbekistan | AY223476 | 81.1 | 85.2 |
| Kosovo/9553/2001 | Yugoslavia | AY675511 | 80.9 | 84.4 |
| 3010 | Congo | DQ019222 | 68.0 | 72.3 |
| Semunya | Uganda | DQ094832 | 67.7 | 71.6 |
| SPU128/81/7 | South Africa | DQ157174 | 81.5 | 85.7 |
| SPU4/81 | South Africa | DQ157175 | 81.6 | 85.7 |
| ROS/HUVLV-100 | Russia | DQ206448 | 81.5 | 84.8 |
| AP92 | Greece | DQ211625 | 70.3 | 72.8 |
| ArD8194 | Senegal | DQ211626 | 67.9 | 72.5 |
| ArD15786 | Senegal | DQ211627 | 67.8 | 72.4 |
| ArD39554 | Mauritania | DQ211628 | 70.5 | 76.5 |
| C-68031 | China | DQ211629 | 87.6 | 89.4 |
| Drosdov | Russia | DQ211630 | 81.2 | 84.0 |
| Kashmanov | Russia | DQ211631 | 81.7 | 84.7 |
| Oman | Oman | DQ211632 | 86.7 | 89.7 |
| SPU97/85 | South Africa | DQ211633 | 91.1 | 91.9 |
| SPU103/87 | South Africa | DQ211634 | 80.8 | 84.4 |
| SPU415/85 | South Africa | DQ211635 | 91.1 | 91.9 |
| Turkey200310849 | Turkey | DQ211636 | 81.7 | 84.3 |
| UG3010 | Congo | DQ211637 | 68.0 | 72.3 |
| Iran-52 | Iran | DQ446215 | 79.6 | 84.9 |
| Iran-53 | Iran | DQ446216 | 80.8 | 84.8 |
| Kosova Hoti | Yugoslavia | EU037902 | 81.7 | 84.6 |
| YL04057 | China | FJ562094 | 69.7 | 72.5 |
| Turkey-Kelkit06 | Turkey | GQ337054 | 81.6 | 83.8 |
| V42/81 | Bulgaria | GU477490 | 80.8 | 83.5 |
| 79121M18 | China | GU477493 | 69.8 | 72.4 |
| Afg09-2990 | Afghanistan | HM452306 | 80.9 | 84.8 |
| Al-Fulah 3-2008 | Sudan | HQ378184 | 80.8 | 85.1 |
| Al-Fulah 9-2008 | Sudan | HQ378185 | 80.8 | 85.1 |
| AB1-2009 | Sudan | HQ378187 | 81.1 | 85.2 |
| NIVA118594 | India | JN572084 | 81.4 | 85.2 |
| NIV112143 | India | JN572085 | 81.4 | 85.2 |
| NIV11703 | India | JN572086 | 81.5 | 85.2 |
| SCT ex Afghanistan | Afghanistan | KC344856 | 83.6 | 86.2 |
| Zahedan | Iran | KC867273 | 80.8 | 84.8 |
| SPU44/08 | South Africa | KJ682805 | 86.1 | 88.7 |
| SPU130/89 | South Africa | KJ682807 | 85.9 | 88.2 |
| SPU45/88 | South Africa | KJ682809 | 86.5 | 87.9 |
| SPU18/88 | South Africa | KJ682810 | 86.1 | 88.4 |
| SPU556/87 | South Africa | KJ682811 | 91.1 | 91.8 |
| SPU431/85 | South Africa | KJ682812 | 90.8 | 91.9 |
| SPU48/90 | South Africa | KJ682813 | 81.3 | 85.2 |
| SPU187/90 | South Africa | KJ682814 | 81.3 | 85.7 |
| Haskovo-V46/13 | Bulgaria | KR092379 | 81.4 | 84.2 |
| 24-R-2012 | Russia | KR814856 | 81.7 | 84.8 |
| 37-R-2013 | Russia | KR814858 | 81.6 | 84.7 |
| 46-ST-2012 | Russia | KR814860 | 81.5 | 85.0 |
| 52-R-2014 | Russia | KR814862 | 81.6 | 84.6 |
| 75-R-2012 | Russia | KR814863 | 81.5 | 84.5 |
| 75-ST-2010 | Russia | KR814864 | 81.7 | 84.9 |
| 3809-ST-2007 | Russia | KR814867 | 81.6 | 84.6 |
| 4495-ST-2008 | Russia | KR814869 | 81.4 | 84.9 |
| 4574-ST-2008 | Russia | KR814870 | 81.7 | 84.8 |
| 4691-ST-2009 | Russia | KR814871 | 81.1 | 84.4 |
| 5180-R-2011 | Russia | KR814874 | 81.6 | 84.7 |
| Oman-199809166 | Oman | KR864901 | 86.7 | 89.5 |
| 1-CR/HU-2015 | Russia | KU161584 | 81.1 | 84.2 |
| Gaib | Tajikistan | KX013445 | 83.1 | 87.2 |
| K128_76 | Kazakhstan | KX013454 | 80.8 | 85.7 |
| K168_40 | Turkmenistan | KX013457 | 80.9 | 85.7 |
| K168_125 | Turkmenistan | KX013460 | 80.8 | 85.5 |
| K229_243 | Russia | KX013466 | 81.2 | 84.7 |
| Min | Russia | KX013481 | 81.2 | 84.7 |
| Saf | Russia | KX013487 | 81.1 | 84.8 |
| Ast133 | Russia | KX056060 | 81.1 | 84.6 |
| YL16070 | China | KY354081 | 69.6 | 72.3 |
| Oman_812956 | Oman | KY362518 | 86.7 | 89.5 |
| UCCR4405 | Russia | KY484029 | 81.2 | 84.1 |
| UCCR4459 | China | KY484032 | 87.5 | 89.4 |
| UCCR4401 | Nigeria | KY484035 | 81.6 | 85.2 |
| UCCR4448 | South Africa | KY484042 | 81.6 | 85.6 |
| UCCR4417 | Uganda | KY484045 | 81.6 | 85.7 |
| Tur_2004_813139 | Turkey | MF511227 | 81.6 | 84.5 |
| Tur_2004_813151 | Turkey | MF511233 | 81.4 | 84.4 |
| SPUD8_81_7_813051 | South Africa | MF511236 | 81.2 | 85.7 |
| 2013_6_180 | Kosovo | MH133278 | 81.3 | 84.7 |
| 2013_14_79 | Kosovo | MH133287 | 81.7 | 84.5 |
| 2014-C-K14-1C | Kosovo | MH133308 | 81.0 | 84.5 |
| 2015-A-K15-1A | Kosovo | MH133311 | 81.2 | 84.4 |
| NIV1311122 | India | MH396643 | 81.2 | 85.2 |
| NIV149247 | India | MH396650 | 81.2 | 85.0 |
| NIV1511559 | India | MH396656 | 80.8 | 85.0 |
| NIV164392 | India | MH396662 | 81.3 | 85.1 |
| 61T/Pakistan | Pakistan | MN135943 | 80.6 | 84.7 |
| 19-H-3116 | India | MN930412 | 80.7 | 85.0 |

**Table S4.** Comparison of nucleotide sequence identity with other CCHFV strains’ L segment

| Strain | County | GenBank accession number | L segment of HANM-18 |
| --- | --- | --- | --- |
|  |  |  | nt (%) |
| Matin | Pakistan | AY422208 | 87.3 |
| TADJ/HU8966 | Tajikistan | AY720893 | 87.4 |
| Baghdad-12 | Iraq | AY947890 | 87.4 |
| IbAr10200 | Nigeria | AY947891 | 86.8 |
| VLV-100 | Russia | AY995166 | 87.4 |
| Semunya | Uganda | DQ076412 | 82.3 |
| 3010 | Congo | DQ099335 | 83.4 |
| AP92 | Greece | DQ211612 | 77.7 |
| ArD8194 | Senegal | DQ211613 | 86.9 |
| ArD15786 | Senegal | DQ211614 | 86.7 |
| ArD39554 | Mauritania | DQ211615 | 86.7 |
| C-68031 | China | DQ211616 | 87.5 |
| Drosdov | Russia | DQ211617 | 87.9 |
| Kashmanov | Russia | DQ211618 | 87.6 |
| Oman | Oman | DQ211619 | 86.5 |
| SPU103/87 | South Africa | DQ211621 | 87.3 |
| SPU415/85 | South Africa | DQ211622 | 87.2 |
| 10849 | Turkey | DQ211623 | 87.9 |
| UG3010 | Congo | DQ211624 | 83.4 |
| Kosova_Hoti | Yugoslavia | EU044832 | 87.9 |
| YL04057 | China | FJ562095 | 93.7 |
| Kelkit06 | Turkey | GQ337055 | 87.8 |
| 79121M18 | China | GU477492 | 93.7 |
| _Afg09-2990 | Afghanistan | HM452307 | 87.2 |
| Al-Fulah_3 | Sudan | HQ378180 | 86.5 |
| Al-Fulah_4 | Sudan | HQ378181 | 86.5 |
| Al-Fulah_9 | Sudan | HQ378182 | 86.5 |
| AB1 | Sudan | HQ378183 | 86.7 |
| NIVA118595 | India | JN572090 | 86.9 |
| NIV112143 | India | JN572091 | 87.0 |
| SCT_ex_Afghanistan | Afghanistan | KC344855 | 87.3 |
| Zahedan | Iran | KC867272 | 87.1 |
| SPU187/90 | South Africa | KJ682795 | 87.3 |
| SPU48/90 | South Africa | KJ682797 | 86.7 |
| SPU431/85 | South Africa | KJ682799 | 87.0 |
| SPU383/87 | South Africa | KJ682801 | 87.0 |
| SPU18/88 | South Africa | KJ682803 | 87.2 |
| SPU497/88 | Namibia | KJ682804 | 87.0 |
| 36-ST | Russia | KR814876 | 87.7 |
| 37-R | Russia | KR814877 | 87.5 |
| 44-ST | Russia | KR814878 | 87.7 |
| 46-ST | Russia | KR814879 | 87.7 |
| 3820-ST | Russia | KR814887 | 87.6 |
| 4691-ST | Russia | KR814890 | 87.7 |
| 1-CR/HU | Russia | KU161582 | 87.5 |
| 1231-CR/TI-2015 | Russia | KU161583 | 87.7 |
| Gaib | Tajikistan | KX013444 | 87.3 |
| Hodzha | Uzbekistan | KX013447 | 87.3 |
| K168_40 | Turkmenistan | KX013456 | 87.3 |
| K229_194 | Russia | KX013462 | 87.5 |
| K229_243 | Russia | KX013465 | 87.5 |
| K315_14 | Russia | KX013468 | 87.4 |
| Mamon | Russia | KX013477 | 87.3 |
| Saf | Russia | KX013486 | 87.6 |
| Ast199 | Russia | KX056050 | 87.3 |
| Ast149 | Russia | KX056053 | 87.6 |
| Ast133 | Russia | KX056059 | 87.7 |
| NIV161064 | India | KY213712 | 86.9 |
| YL16070 | China | KY354082 | 93.7 |
| UCCR4413 | Pakistan | KY484039 | 87.2 |
| UCCR4417 | Uganda | KY484043 | 87.2 |
| Kalmykia_Shch_1 | Russia | KY982864 | 87.4 |
| Tur_2004_813048 | Turkey | MF511190 | 87.9 |
| Tur_2004_813135 | Turkey | MF511191 | 87.7 |
| Tur_2004_813137 | Turkey | MF511192 | 87.8 |
| SPU41_84_813060 | South Africa | MF511206 | 86.5 |
| 2013_6_180 | Kosovo | MH133279 | 88.1 |
| 2013_14_79 | Kosovo | MH133288 | 88.0 |
| 2013_19_289 | Kosovo | MH133297 | 88.0 |
| 2013_25_380 | Kosovo | MH133303 | 87.9 |
| A-K14-1A | Kosovo | MH133306 | 87.7 |
| A-K15-1A | Kosovo | MH133312 | 87.6 |
| NIV1733666 | India | MH396673 | 87.0 |
| YL16204 | China | MH688499 | 93.7 |
| MCL-18-T-305 | India | MK330136 | 87.2 |
| 19-H-3068 | India | MN930403 | 87.1 |
| 19-H-2856 | India | MN930405 | 87.2 |
| 19-H-3092 | India | MN930408 | 87.2 |
| 19-T-1989 | India | MN930410 | 87.1 |

